# Mouse Adapted SARS-CoV-2 Model Induces “Long-COVID” Neuropathology in BALB/c Mice

**DOI:** 10.1101/2023.03.18.533204

**Authors:** Timothy E. Gressett, Sarah R. Leist, Saifudeen Ismael, Grant Talkington, Kenneth H. Dinnon, Ralph S. Baric, Gregory Bix

**Affiliations:** Department of Neurosurgery, Clinical Neuroscience Research Center, Tulane University School of Medicine, New Orleans, LA 70112, USA; Tulane Brain Institute, Tulane University, New Orleans, LA 70112, USA; Department of Epidemiology, University of North Carolina at Chapel Hill, Chapel Hill, North Carolina, USA; Department of Microbiology and Immunology, University of North Carolina at Chapel Hill, Chapel Hill, NC, USA; Department of Neurology, Tulane University School of Medicine, New Orleans, LA 70112, USA; Department of Microbiology and Immunology, Tulane University School of Medicine, New Orleans, LA 70112, USA

**Author notes:** Authors have equal contribution to this study (Timothy E. Gressett, Sarah R. Leist, Saifudeen Ismael). **Correspondence: Gregory Bix, M.D., Ph.D., F.A.H.A.**, Tulane University School of Medicine, Center for Clinical Neurosciences, 131 S. Robertson, Ste 1300, Room 1349, New Orleans, LA 70112.

**Keywords:** SARS-CoV-2, MA10, Long COVID, Neuropathology, NeuN, Iba-1, Hippocampus

## Abstract

The novel coronavirus SARS-CoV-2 has caused significant global morbidity and mortality and continues to burden patients with persisting neurological dysfunction. COVID-19 survivors develop debilitating symptoms to include neuro-psychological dysfunction, termed “Long COVID”, which can cause significant reduction of quality of life. Despite vigorous model development, the possible cause of these symptoms and the underlying pathophysiology of this devastating disease remains elusive. Mouse adapted (MA10) SARS-CoV-2 is a novel mouse-based model of COVID-19 which simulates the clinical symptoms of respiratory distress associated with SARS-CoV-2 infection in mice. In this study, we evaluated the long-term effects of MA10 infection on brain pathology and neuroinflammation. 10-week and 1-year old female BALB/cAnNHsd mice were infected intranasally with 10^4^ plaque-forming units (PFU) and 10^3^ PFU of SARS-CoV-2 MA10, respectively, and the brain was examined 60 days post-infection (dpi). Immunohistochemical analysis showed a decrease in the neuronal nuclear protein NeuN and an increase in Iba-1 positive amoeboid microglia in the hippocampus after MA10 infection, indicating long-term neurological changes in a brain area which is critical for long-term memory consolidation and processing. Importantly, these changes were seen in 40-50% of infected mice, which correlates to prevalence of LC seen clinically. Our data shows for the first time that MA10 infection induces neuropathological outcomes several weeks after infection at similar rates of observed clinical prevalence of “Long COVID”. These observations strengthen the MA10 model as a viable model for study of the long-term effects of SARS-CoV-2 in humans. Establishing the viability of this model is a key step towards the rapid development of novel therapeutic strategies to ameliorate neuroinflammation and restore brain function in those suffering from the persistent cognitive dysfunction of “Long-COVID”.

## 1. Introduction

The World Health Organization defines “Long-COVID” (LC) as a condition that occurs in individuals with a history of probable or confirmed SARS-CoV-2 infection, usually three months from the onset of COVID-19, with the two most frequently reported symptoms beyond six months as both fatigue and cognitive dysfunction [1, 2]. Unfortunately, up to 40-50% of individuals that have been infected with SARS-CoV-2 will go on to experience LC symptoms, with persistent neurological symptoms such as fatigue and memory impairment [3–5]. Currently, no animal models for LC exist which fully recapitulate the disease [6]. Thus, our ability to understand the underlying pathophysiology of this debilitating condition is significantly constrained [7], limiting our ability to develop effective therapeutic strategies.

Previously, two models have been utilized to study the acute effects of SARS-CoV-2 in laboratory mice. The first model is an adenoviral (Ad5) mediated vector model [8] which allows for exogenous delivery of human ACE2 (hACE2) under the CMV promoter into pulmonary epithelial cells. These Ad5-hACE2 transduced mice show evidence of viral infection to include perivascular and interstitial inflammatory cell infiltrates, necrosis, edema, and systemic effects as seen in human SARS-CoV-2 infection. The other model used is a more contemporary transgenic model (K18-hACE2) which expresses hACE2 under the human keratin 18 (K18) promoter in airway epithelia where SARS-CoV-2 infections typically begin [9]. These mice also show evidence of viral infection in the lungs with the addition of dose-dependent damage to systemic organs in a manner consistent with clinical observations in postmortem samples from COVID-19 patients [10]. Unfortunately, the use of these models to study the long-term sequalae of SARS-CoV-2 is problematic [11, 12]. Although both models are able to show pathophysiological outcomes inherent to SARS-CoV-2 infection [13], they are inherently confounded by the use of a human receptor in a murine background which is not being expressed by its native mouse promoter. Therefore, the accuracy of these models to study the long-term neurological effects of SARS-CoV-2 infection and their clinical translational value, remain uncertain.

Here we present evidence that a novel mouse-adapted SARS-CoV-2 (MA10) model which has been previously validated to recapitulate the major clinical manifestations and acute respiratory distress observed in humans [14] may provide a way to study the long-term neurological effects of SARS-CoV-2 infection. Importantly, the MA10 strain has been extensively studied [14, 15] in standard laboratory BALB/c mice which show pathophysiological responses [16]. We are the first, however, to evaluate the long-term neurological impact of SARS-COV-2 MA10 infection in these standard laboratory BALB/c mice. As currently available models may not be adequate for long-term neuropathological study of SARS-CoV-2 [17–19], the use of a mouse-adapted strain of SARS-CoV-2 may allow for better translational research that is more relevant to native physiology which uses the native mouse ACE2 receptor under its operant promoter. In this study, we investigated the long-term neurological effects of SARS-CoV2 “MA10” infection as a potential tool in establishing a clinically relevant model for the development of novel therapeutic strategies for “Long-COVID”.

## 2. Methods

### 2.1. SARS-CoV-2 Infection

10 week and 1-year old female BALB/cAnNHsd (“BALB/c” strain 047) mice were purchased from Envigo and housed at the University of North Carolina at Chapel Hill then infected with SARS-CoV-2 MA10 under biosafety level 3 (BSL3) conditions as described previously [14]. All animal work was approved by appropriate institutional animal care and use committees. Briefly, mice were inoculated with either mock (saline) or MA10 strain of severe acute respiratory syndrome coronavirus 2 (SARS-CoV-2, BEI resource, NR-55329) via intranasal administration at a dose of 1×10^4^ plaque-forming unit (PFU) or 1×10^3^ PFU, respectively. After 60 days postinfection (dpi), mice were euthanized by isoflurane overdose and brains were collected at the University of North Carolina and shipped to Tulane University School of Medicine for paraffin embedding, sectioning, and staining for immunofluorescent analysis.

### 2.2. SARS-CoV-2 Immunofluorescence Analysis

Immunofluorescent preparation and analysis were conducted as previously described [20]. Briefly, brains were embedded in paraffin and sectioned at 5 μm thickness, deparaffinized in xylene, and rehydrated through a series of ethanol washes followed by heat-induced antigen retrieval with high pH antigen unmasking Tris-EDTA solution. Slides were then washed with PBS with 0.3% Triton X-100 and blocked with 5% normal goat serum for 1h at room temperature. The brain sections were incubated with primary antibody (Rabbit anti-Iba1, at 1:500, Cat# 019-19741, and anti-NeuN (Cat#ab128886, 1:250) at 4°C overnight. Slides were then washed and appropriate secondary antibody (Jackson Immunoresearch, Cat#615545-214, 1:500) tagged with Alexa Fluor fluorochromes in normal goat serum was used for 1h at room temperature. Mounting media with DAPI was used to label the nuclei after washing with PBS. Slides were then scanned with a digital slide scanner (Zeiss Axio Scan; Zeiss, White Plains, NY) at 20X magnification and analyzed using QuPath software (version 0.3.0, University of Edinburgh) [21]. Both NeuN and Iba-1 immunofluorescence were quantified manually after regional segmentation of the hippocampus using the Franklin and Paxinos mouse atlas [22]. Counting of NeuN in each region of the hippocampus was defined by complete circumferential signal intensity of each cell. Scoring of Iba-1 positive microglia in the hippocampus was done via cytoplasm analysis and defined as G1 (0-30%), G2 (30-60%), and G3 (60-100%) as relative percentages of signal intensity based on previously characterized ameboid-like Iba-1 positive microglia phenotypes which exhibits a hypertrophic cell body correlating to an activated state [23, 24]. All scoring was done blinded.

### 2.3. Statistics and Hypothesis

Statistical tests were performed using GraphPad Prism, 9.3.1 version (GraphPad Software, San Diego, CA). Data are presented as mean ± SEM. Significant differences were designated using omnibus one-way ANOVA and, when significant, followed-up with two-group planned comparisons selected *a priori* to probe specific hypothesis-driven questions. Statistical significance was taken at the *p* < 0.05 level. In this study we tested the hypothesis that MA10 infected mice would show neuropathological changes with the same clinical symptom prevalence as reported in those diagnosed with “Long COVID”.

## 3. Results

### 3.1. Neuronal marker NeuN is decreased in the hippocampus of 60dpi mice after MA10 infection

Immunofluorescence analysis with neuronal nuclear antigen marker NeuN showed a decrease in NeuN positive cells between both old and young MA10 infected mice at 60 dpi in the hippocampus (Fig 1B) in comparison with mock animals. When both young and old groups were combined the trend of decreased NeuN positive cells increased towards significance (Fig 1C). Interestingly, when infected animals are combined irrespective of their age, a subset of MA10-infected mice (~45%, or n=5) showed a significant decrease (*p<0.05, two-tailed unpaired Student’s t-test) in NeuN positive cells count (Fig 1A and 1D). This approximates what is observed clinically in terms of “Long COVID” symptom prevalence [3–5].

**Fig. 1.**
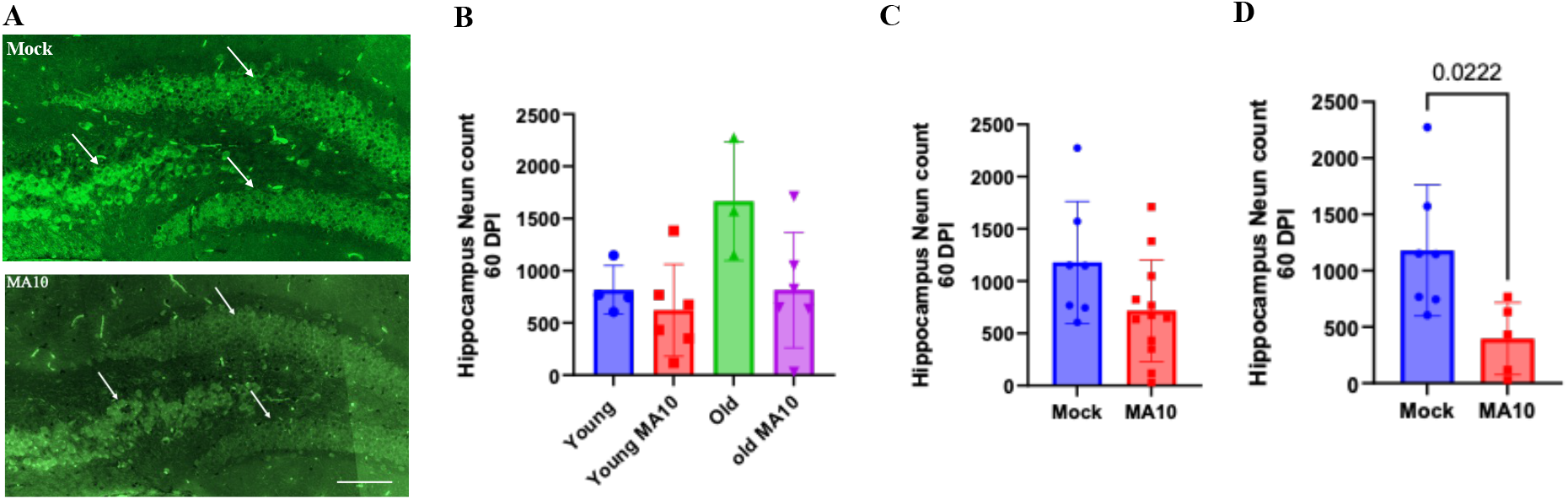
MA10 infection decreases neuronal NeuN density in the hippocampus of BALB/cAnNHsd mice at 60 days post infection. A. Hippocampal images of Iba-1 positive microglial cells (green), 60 days after intranasal inoculation with saline (mock) or SARS-CoV-2 MA10 (1×10^3^ PFU) in 1-year-old female mice. B. Quantification of NeuN positive cells between young (10-week) and old (1-year) demonstrating a marginal decrease in NeuN positive cells. C. Combination of both young and old subgroups showed a more pronounced trend of decreasing NeuN positive cells. D. Quantification of NeuN positive cells as in A demonstrating significant decrease (*p<0.05, two-tailed unpaired Student’s t-test) in NeuN positive cells in subset (~45%) of MA10-infected mice which showed pronounced pathology. Data are presented as mean ± SEM. Saline n=8; MA10 n=5 scale bar = 25 μm.

### 3.2. Microglia marker Iba-1 is increased in the hippocampus of 60dpi mice after MA10 infection

Staining with microglial-specific calcium binding protein marker Iba-1 showed an increase in amoeboid microglia cells between old and young MA10 infected mice at 60 dpi in the hippocampus (Fig 2B). When both young and old groups were combined, the trend of Iba-1 positive amoeboid microglia cells in the G3 cytoplasm increased towards significance (Fig 2C). Strikingly, significance between groups (*p<0.05, two-tailed unpaired Student’s t-test) of increased Iba-1 positive amoeboid microglia cells appeared in the same subset (~45%) of MA10-infected mice (Fig 2A and 1D) as previously quantified for NeuN.

**Fig. 2.**
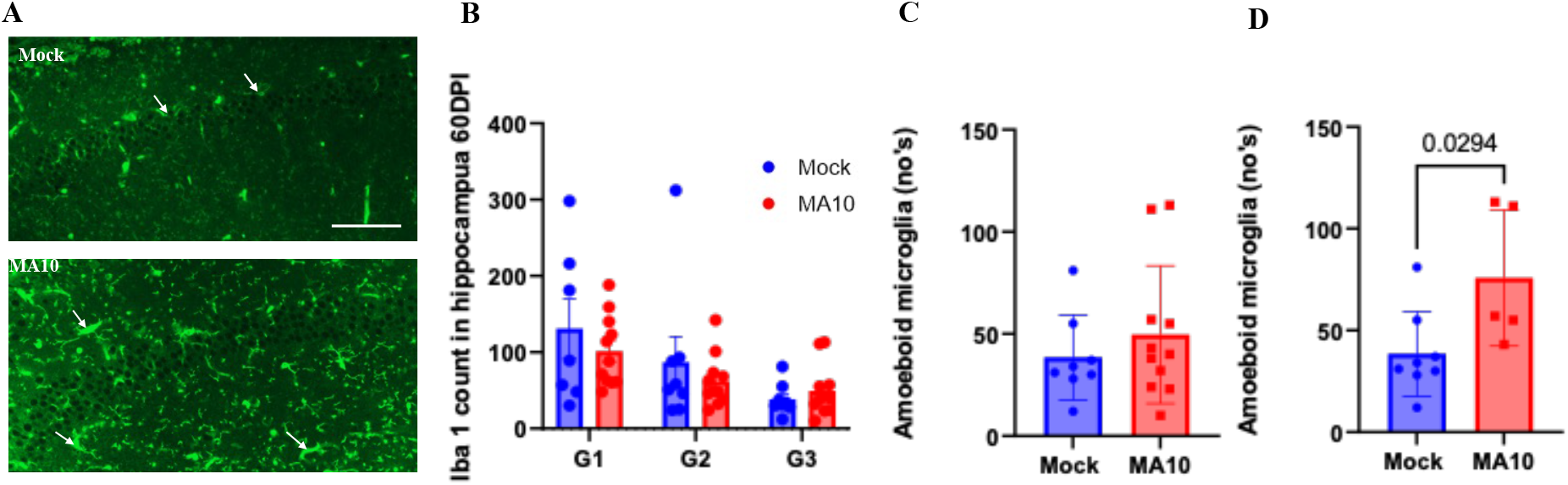
MA10 infection increases Iba-1 positive amoeboid microglial cells in the hippocampus of BALB/cAnNHsd mice at 60 days post infection. A. Hippocampus images of Iba-1 positive microglial cells (green) 60 days after intranasal inoculation with saline (mock) or SARS-CoV-2 MA10 (1×10^3^ PFU) in 1-year-old female mice. B. Quantification of Iba-1 positive microglial cells as in A demonstrating a marginal increase in Iba-1 positive amoeboid microglia. C. Combination of both young and old subgroups showing an increasing trend of increasing Iba-1 positive cells. D. Quantification of Iba-1 positive microglial cells as in A demonstrating a significant increase (*p<0.05, two-tailed unpaired Student’s t-test) in Iba-1 positive cells in subset (~45%) of MA10-infected mice which showed pronounced pathology. G1 (0-30%), G2 (30-60%), and G3 (60-100%) as percentages of signal intensity based on previously characterized ameboid-like Iba-1 positive microglia phenotypes which exhibits a hypertrophic cell body correlating to an activated state. Data are presented as mean ± SEM. Mock n=8; MA10 n=5, scale bar = 25 μm.

## 4. Discussion

In this study, we sought to characterize the long-term neuropathological changes induced after infection with a novel mouse adapted SARS-CoV-2 (MA10) in standard laboratory mice. As the full extent of neuropathological changes induced by SARS-CoV-2 are currently unknown and current models may not accurately recapitulate the disease adequately, we sought to examine key markers in the brain relevant to neuronal and immune function after viral inoculation relevant to the disease course and severity in mice as compared to humans [14, 25].

This study demonstrated that SARS-CoV-2 MA10 infection can cause a decrease in NeuN positive neurons and an increase in pathological Iba-1 positive microglia at 60 days post infection. Importantly, this effect was only seen in a subset (~45%) of mice. Previously, we conducted studies in non-human primates to model the neurological effects of SARS-Cov-2 in rhesus macaques and African green monkeys, where we found increased blood-brain barrier permeability as well as the presence of cerebral microhemorrhages and microinfarcts consistent with those found in imaging reports from critically ill human COVID-19 patients [26]. This agrees with clinical observations recently detailed in a comprehensive systematic review that approximately half of those infected with SARS-CoV-2 will experience lingering symptoms [27], and validates the ~45% of neuropathology seen in our MA10 mouse model studies.

Caution must be taken in interpreting our results. The lack of NeuN in the hippocampus may indicate transcriptional changes and not a complete loss of neuronal function between groups, as NeuN is a marker for neuronal nuclear protein [28]. Additionally, our analysis of Iba-1 is based on cytoplasmic density as an indicator of pathologically activated amoeboid microglia [23, 24], which, although we believe is the most reliable measurement to assess actin-bundling activity and thus the membrane ruffling and phagocytotic state in activated microglia [29], other ways to interpret our analysis may produce a different conclusion on pathological activation. Lastly, as it is known that Iba-1 marks both microglia and infiltrating macrophages [30], additional studies may be necessary to determine the exact immune response caused by MA10 infection. Taken together, however, the consistent pathological findings across different markers after subset analysis strongly resemble observed clinical features of “Long-COVID”, validating SARS-CoV-2 MA10 as a plausible model for studying this disease and generating novel treatments for disease severity and progression. Future directions of this work may seek to establish a more rigorous time-series of observable neuropathology and link this neuropathology to behavioral studies to examine its effect of cognitive dysfunction. Although it has been shown that SARS-CoV-2 spike protein alone is capable of inducing TLR4-mediated long-term cognitive dysfunction in mice [31], this result has yet to be repeated using live virus.

In summary, our findings indicate that the SARS-CoV-2 (MA10) infected mouse model induces neuropathology in standard laboratory mice which persists up to 60 days post infection. This may make it an ideal candidate for the study of long-term neuropathological changes induced by SARS-CoV-2 infection and warrants further study as a suitable model for “Long COVID” and subsequent therapeutic testing. A distinct advantage of this model is that it preserves native-host ACE2 expression and does not rely on the insertion of hACE2 receptor under a different promoter, as previous models have used. Future directions of this work warrant therapeutic targeting to ameliorate infectivity and neuropathology associated with SARS-CoV-2, which remains a primary focus of our group. Being a preliminary study, our evaluations are limited to a few markers of long-term neuropathological changes and require further studies to determine the exact mechanisms of MA10-associated neuropathology. Furthermore, long-term experimental studies are needed to assess functional and behavioral outcomes to assess for cognitive dysfunction and fatigue which are predominant clinical features of “Long COVID”. Lastly, additional studies with this novel MA10 model may allow for a variety of therapeutic testing with rapid turnover using standard laboratory mice, which may be used to study long-term neuropathological consequences of SARS-CoV-2 and alleviate corresponding clinical sequalae of “Long COVID” in humans.

## Author Contributions

Conceptualization, GB.; methodology, T.E.G., S.R.L., S.I., G.T., K.H.D., R.S.B., and G.B; validation, T.E.G., S.I.; formal analysis, T.E.G., S.I., G.T., writing—original draft preparation, T.E.G., S.I., and G.T.; writing—review and editing, T.E.G., S.R.L., S.I., G.T., K.H.D., R.S.B. and G.B.; resources and funding acquisition, R.S.B. and G.B. All authors have read and agreed to the published version of the manuscript.

## Data Availability Statement

All relevant data is included in this article. SARS-CoV-2 MA10 is available from BEI resources.

## Acknowledgments

We would like to extend a special thanks Dr. Ralph Baric at the University of North Carolina at Chapel Hill for his continued support and insightful feedback throughout the completion of the project.

